# Opposite-sex associations are linked with annual fitness, but sociality is stable over lifetime

**DOI:** 10.1101/2022.01.04.474937

**Authors:** Jamie Dunning, Terry Burke, Alex Hoi Hang Chan, Heung Ying Janet Chik, Tim Evans, Julia Schroeder

## Abstract

Animal sociality, an individual’s propensity to associate with others, has fitness consequences through mate choice, for example, directly, by increasing the pool of prospective partners, and indirectly through increased survival, and individuals benefit from both. Annually, fitness consequences are realised through increased mating success and subsequent fecundity. However, it remains unknown whether these consequences translate to life-time fitness. Here, we quantified social associations and their link to fitness annually and over lifetime, using a multi-generational, genetic pedigree. We used social network analysis to calculate variables representing different aspects of an individual’s sociality. Sociality showed high within-individual repeatability. We found that birds with more opposite-sex associates had higher annual fitness than those with fewer, but this did not translate to lifetime fitness. Instead, for lifetime fitness, we found evidence for stabilizing selection on opposite-sex sociality, and sociality in general, suggesting that reported benefits are only short-lived in a wild population, and that selection favours an average sociality.

## Background

Some individuals are consistently more sociable than others, demonstrated by within-individual repeatability of social traits across vertebrate groups (Aplin et al. 2015; Thys et al. 2017; Dimitriadou, Croft and Darden 2019; Plaza et al. 2019; Beck, Valcu, and Kempenaers 2020; Proops et al. 2021; Strickland et al. 2021). This variation in individual sociality is positively linked with fitness in some taxa (Silk 2007; Silk et al. 2009) and is therefore expected to be subject to selection (Krause and Ruxton 2002). Fitness is a relative measure of an individual’s genetic contribution to the population in the next generation, and thus, can only be represented comprehensively and precisely by measures of traits spanning an organism’s lifetime (Endler 1986; Blankenhorn 2010; Reid et al. 2019; Moiron, Charmantier and Bouwhuis 2022). A comprehensive definition of fitness is fundamental to understand the evolutionary pressures that shape variation in sociality. In practice, however, many studies must rely on fitness correlates instead (e.g., number of broods, or survival, instead of genetic contribution). For example, in mammal societies, both variation in within- and between-sex affiliations (Archie et al. 2014) have been linked to lifetime fitness correlates, survival and longevity (Cameron, Setsaas and Linklater 2009; Silk, Alberts and Altman 2003; Silk et al. 2010; Stanton and Mann 2012, but also see Thompson and Cords 2018). Whereas, in birds, the subject of this study, the use of fitness correlates (eggs laid, chicks fledged, within-year survival etc.) are frequent over more precise fitness measures (Moiron, Charmantier and Bouwhuis 2022), that require intensive field work over a long period of time.

Although some benefits are linked with sociality during the breeding period most of these tend to be short-term and contextual (Bebbington et al. 2017; Riehl and Strong 2018). Instead, benefits associated with reproduction are often linked with non-breeding sociality (Firth and Sheldon 2016; Kohn 2017; Maldonado-Chaparro et al. 2018; Beck, Farine, and Kempenaers 2020), when group cohesion is stronger (Silk et al. 2014; Kurvers 2020). Sociality may influence fitness in different ways, through benefits to reproductive success or increased survival, and so, the mechanism of selection acting on social traits may also differ. For example, communal foraging between socially associated individuals during the non-breeding period facilitates resource information transfer (Aplin et al. 2012; Firth, Sheldon and Farine 2016; Hillemann et al. 2020) and reduces predation risk (Cresswell 1994; Cresswell and Quinn 2011; Sorato et al. 2012), increasing survival. However, these benefits may also incur costs associated with competition for resources and mate choice (Birkhead and Biggins 1987; Le Galliard et al. 2005; Forstmeier et al. 2011; Mayer and Pasinelli 2013; Grant and Grant 2019; but also see Lea et al. 2010). Sociality may also benefit individuals who hold more central social network positions or have access to opposite-sex associates, through enhansed mate choice (McDonald 2007; Oh and Badyaev 2010; Firth et al. 2018; Beck, Farine, and Kempenaers 2021). Although the association between annual fitness correlates and non-breeding sociality has been well described, testing how selection acts on social traits requires lifetime fitness measures, and remains unresolved.

With the recent development of tools to construct and analyse social networks (Wey et al. 2008; Farine and Whitehead 2015) the study of sociality has become popular among behavioural ecologists. Yet, to describe the association between sociality and fitness any potential study must first overcome two problems: (1) A social association must be clearly defined relative to the behavior of the study system (Figure 1; Psorakis et al. 2012, 2015), and (2) to study the evolution of social behaviour, precise measures of individual fitness must be quantified to test for correlation with a social trait – selection. Although annual fitness correlates are widely used, lifetime fitness is more precise and without the stochasticity of annual measures (Dobson, Murie and Viblanc 2020; Alif et al. 2022) and thus, can better describe selection pressure acting on a trait (Endler 1986; Blankenhorn 2010; Reid et al. 2019). However, lifetime fitness requires wild animals to be monitored throughout their whole lives, and all breeding attempts, and the fates of their offspring, must be recorded to determine recruitment. All of these require a multi-generational, genetic pedigree (Kruuk 2004; Korsten et al. 2013).

**Figure 1.**
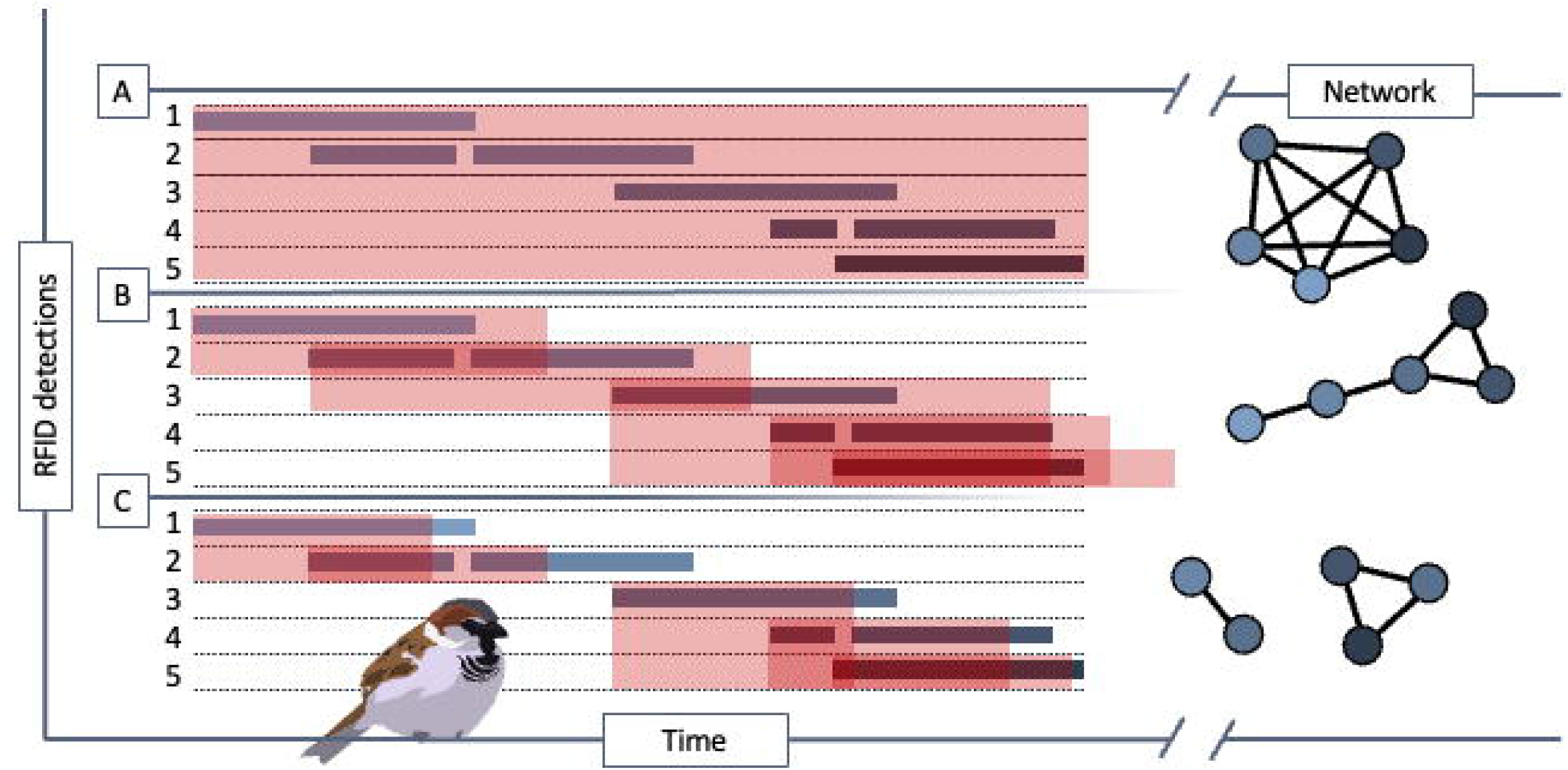
Three versions of a simulated event (A, B and C) show the interval over which five individuals (1–5, black/grey bars) spent at a resource over time (_*t*_), and the derived social networks from each approach: A = gambit of the group, which links all individuals in a discrete group equally; B = time-window overlap (by △_*t*_), which links individuals who overlap at a resource; and C = arrival time (developed for this study), which links individuals who arrive together to a resource. Red boxes denote the time period during which individuals are considered to be associated, and overlaps represent an association: A, all individuals within a group; B, where they are physically present at the same time (red box), or shortly after they depart to account for birds which were present, but not currently being recorded by the antenna, in that case, over-lapping by △_*t*_ (red over-hanging box, typically a few seconds); or, C, where they arrive within △_t_ of each other, but the subsequent time spent at a feeder is irrelevant. However, note that the function of △_t_ differs between B and C; Where in B, △_t_ functions to detect when birds are in the same place but where one (or more) are not currently being detected by the antenna, in C the function is to link all individuals which arrive together, while ignoring those already present at the resource, which has the potential to link two separate groups in A and B. In the case of C, an additional interval (△_i_) is required to define when birds have left the resource, after which they can be recorded as arriving again.

Our study system, an island population of house sparrows *Passer domesticus* (hereafter, sparrow/s) where we monitor all individuals from birth to death, without capture bias (Simons et al. 2015), overcomes both problems. (1) We have sociality data from birds that are electronically registered visiting a feeder. Social centrality measures are repeatable across different timescales and contexts in this and other populations (Plaza et al. 2019). (2) We have lifetime recruitment data available, from a multi-generational genetic pedigree that, because our population is closed, meaningly there is no movement of sparrows to or from the island, and that our study covers all sparrows on the island, we can use to compute precise annual and lifetime fitness estimates (Schroeder et al. 2015; Alif et al. 2022).

We tested predictions based on arguments presented above to understand the potential for selection on sociality: (1) We confirmed that the social traits we measured were meaningful by testing for individual repeatability of sociality; (2) We tested the prediction that non-breeding sociality has fitness benefits, either driven by reproductive success through opposite-sex association or through increased survival through network centrality measures and is subject to selection.

## Methods

### Study system

We used data from the Lundy sparrow system, a long-term study based on the island of Lundy (51.11N, 4.40W), ≈19 km off North Devon, UK, in the Bristol Channel. The sparrows on Lundy breed in nest boxes, sited in groups around the only village on the island. The island is rodent-free and therefore the sparrows have no predators but for the occasional vagrant raptor. House sparrows are a model organism in behavioral ecology and evolution, and much is known about their biology, physiology and life-history (Andersson 1994; Sánchez-Tójar et al. 2018). House sparrow are socially monogamous, but 25% of broods show they can be genetically promiscuous (Schroeder et al. 2016). On Lundy, they have on average 2-3 broods of 4-5 eggs per breeding season (Westneat et al. 2014). The sex ratio is stable, and the mean lifespan of recruits is three years (Alif et al. 2022). Although sparrows are territorial during the breeding season, during the non-breeding period they form gregarious groups that forage communally for seed and at supplementary bird feeders (Summer-Smith 1963), both of which are available year-round on Lundy.

Most sparrows were first captured, and tissue sampled in nest boxes at their natal site during the breeding season (April to August) or using mist nets during the post-fledging period (Schroeder et al. 2011; Girndt et al. 2019). Tissue samples were either blood or mouth swabs and were stored in ethanol and refrigerated at 3°C prior to analysis. We genotyped sparrow DNA at <22 microsatellite loci suitable for parentage assignment in sparrows (Dawson et al. 2012). Using the genetic data, we assembled a near-complete genetic pedigree (Schroeder et al. 2015, 2016), which at the time of writing spans twenty years, 2000–2019, and 8,379 individuals. We fitted all sparrows with a unique combination of a coded metal ring and three coloured leg rings. We also provided each sparrow with a subcutaneous Passive Identification Transponder (PIT tag; TROVANID100: 11.5 × 2.1 mm and 0.1 g), under the skin of the breast, which we have previously shown have no detrimental effect on subsequent fitness (for details see Schroeder et al. 2011). These tagged individuals were then recorded when they visited a custom made 19.8cm x 19.8cm Radio Frequency Identification antenna (RFID; DorsetID) mounted on a seed reservoir (for photo see Sánchez-Tójar et al. 2017; Brandl et al. 2019), positioned centrally within our study site. The feeder was open access, and explicitly not limited to a single bird feeding at one time, as is the case at hanging bird feeders (Youngblood 2019; Beck, Farine and Kempenaers 2020). Our feeder recorded visiting birds every day that the island generators were running (6am – midnight, seven days a week).

### Social centrality measures

To quantify the sociality of individual sparrows we calculated measures of social centrality (hereafter centrality measures) using presence data from the RFID antenna, collected during the non-breeding periods, November–January, of 2015/16 (15 weeks) and 2016/17 (13 weeks, hereafter referred to events). An association between birds observed from these data can reflect either, individuals that choose to maintain some social cohesion, which we consider to be a non-random association, or individuals without pre-established social cohesion who coincide in time and space, which we consider random mixing. These data record the presence of birds without distinguishing between these two types of association. Further, dyadic interactions can also be aggressive interactions, and our data cannot exclude these cases. Although in sparrows’, dominance hierarchies are typically linear, there is no evidence for a correlation with reproductive fitness (Sánchez-Tójar 2018).

The ‘gambit of the group’ is a common approach used to identify discrete groups among all associating individuals (Whitehead and Dufault 1999; Figure 1A). However, given the gregarious nature of sparrows and the high activity at our feeder, at which non-discrete groups of sparrows accumulated, the gambit of the group approach overestimated associations between individuals (Figure 1A; also see Ferreira et al. 2020). One solution to this would be to define a non-random association where two individuals overlapped by a defined period (△_t_) at our bird feeder. However, in our system, again due to the near constant visitations, this resulted in linear network structures, e.g., linking the first bird to the second, then the second to the third, and so on (Figure 1B). To account for the social behaviour of sparrows, we derived a method to infer non-random associations that assumed that non-random social associations are established before they attend the feeder (suggested by Summer-Smith 1963). We therefore defined an association as two individuals that arrived to feed within 150 seconds (△_*t*_) of each other. Here an arrival is defined as the (re)appearance of the individual at the feeder after being absent for a period of minimally >300 seconds (△_*t*_). We defined that △_*t*_ = 150 seconds was sufficient to detect and link individuals who arrive together in a group (see Figure 1C), and the resulting data better sampled non-random associations between individuals in our system, from watching sparrows in the field and on pre-recorded footage, arriving at our feeder (Plaza et al., 2019).

From the resulting association matrices of the two events, 2015/16 and 2016/17, we built a series of weighted, non-directional, social networks (hereafter, network/s), where the vertices represent individual sparrows and interconnecting edges their associations. First, we built individual networks for each of the 15 weeks in 2015/16, and 13 weeks in 2016/17, to estimate within-individual repeatability in centrality network metrics. This was to validate these centrality measures against the individual repeatability already demonstrated in this system using both RFID and video data (Plaza et al. 2019). Then, we built a network for each event to extract non-breeding sociality. Finally, we also built two bipartite networks from each event (sub-graphs), which only considered association strength between opposite-sex individuals.

From the first non-breeding networks we extracted three centrality measures representing different aspects of sociality for each individual using the ‘iGraph’ R package (Csardi and Nepusz 2006): We selected centrality measures to reflect aspects of an individual’s social preference, following similar studies on sociality (McDonald 2007; Farine and Whitehead 2015; Beck, Farine and Kempenaers 2021). Degree represents the number of associates, and may impact fitness through enhanced mate choice, where individuals position themselves alongside others of lesser quality (Oh and Badyaev 2010;), or where same-sex associates benefit reproduction in cooperative breeding species (Bebbington et al. 2017; Riehl and Strong 2018). Strength represents the quality of those relationships and may influence the structure and behaviour of reproductive communities (Firth and Sheldon 2016; Culina, Firth and Hinde 2020). We calculated strength following (Farine 2013), using the sum of dyadic Simple Ratio Indices (the association probability between a dyad, from 0, never associated, to 1, always associated), which we transformed to give a measure of net association quality (Eq. 1, where S denotes strength, d_(i)_ and sum SRI_(i)_ the degree and SRI of a given individual respectively and N(V) = the number of vertices in a network)

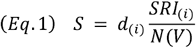

We used eigenvector centrality (following McDonald 2007, hereafter centrality) to quantify the influence of an individual to all others within the network (Newman 2004; Oh and Badyaev 2010); Finally, we extracted opposite-sex degree from the two bipartite sub-graphs, the number of opposite-sex associations, which we used to represent an individual’s pool of potential reproductive partners, as a fourth measure of sociality (following Beck, Farine and Kempenaers 2021). Given the high density of sparrows visiting the feeder, and the frequency at which those birds were detected we did not threshold our networks (using only a sample of the birds attending an antenna, following Farine and Whitehead 2015) to maintain network structure, although only birds who arrived in a dyad (where degree > 0) were included in our networks.

### Fitness measures

For each of the sparrows that survived to the following breeding period we used the genetic pedigree to calculate two fitness measures, and for each of these, we calculated one annual measure, and one across the lifetime of the sparrow. Both fitness measures are based on the number of recruits, and we have shown that they correlate well with reproductive value, thus represent fitness in an evolutionarily meaningful way (Alif et al. 2022). We defined recruits as offspring that survived and produced genetic offspring themselves.

#### Recruits

For the number of annual recruits, we summed individual recruits within the breeding year following the social events. We then again summed individual recruits across a lifetime, or up to 2020 as a measure of lifetime recruits. Note that the latter category only contained five sparrows that were still alive at the point of census, and as such, our recruitment data can be considered near complete. We excluded birds which did not survive to breed, and yearlings which had zero recruits.

#### De-lifed fitness

As a second fitness measure, we used de-lifed fitness (*p*^*ti*^ ; Eq. 2), which estimates an individual’s genetic contribution to the population (Coulson et al. 2006). De-lifed fitness is a retrospective measure of realized fitness, relative to the population each year, calculated by removing (de-lifing) an individual and its offspring from the pedigree and recalculating the resulting change in population growth.

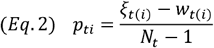

Here, *p*_*ti*_ is the contribution of individual (_*i*_) to population growth during a specific period *t*. Further, *ξ*_*t*(*i*)_ is a measure of individual performance, here the number of surviving offspring of individual *i* at the end of the breeding period *t*. We added a value of one if the individual *i* itself survived to the next breeding period *t* + 1. The population size at time *t* is *N*_*t*_ at the beginning of each breeding cycle (here April). To estimate the individual’s contribution to population growth, we use *w*_*t*_, which represents the ratio of the population size at *t* + 1 to the population size at *t*. This de-lifed fitness *p*_*ti*_ is an annual value per individual, and we calculated it for all birds which produced at least one recruit. We then also summed *p*_*ti*_, within individuals as a lifetime de-lifed fitness measure, *p*_*i*_.

### Individual repeatability in sociality

First, we validated our centrality measures by confirming that they were repeatable within-individuals between weeks. We modelled degree, strength, centrality and opposite-sex degree respectively, as response variables each in four Gaussian linear mixed models against the intercept, and with bird identity modelled as a random effect to compensate for repeat identities between years. We then divided the variance explained by bird identity by the total phenotypic variance of the trait to quantify repeatability (see Nakagawa and Schielzeth 2010). We ran repeatability models using package MCMCglmm default parameters and priors – the models converged robustly and reliably.

### Selection on sociality – annual and lifetime fitness

We quantified the association between centrality measures and fitness. As all four centrality measures are inherently correlated, we modeled each separately to avoid collinearity (Webster, Schneider and Vander Wal 2020). For each centrality measure, we ran two models, with annual recruits and annual de-lifed fitness as the response variables. In the models with annual numbers of recruits as response, we assumed a Poisson error distribution with a log link function, and in the models explaining de-lifed fitness we assumed a Gaussian error distribution. We mean-centered all centrality measures within each year, eliminating between-year differences, and modelled them as fixed covariates. We also added each sociality variable as a quadratic effect to test for stabilizing or disruptive selection where averages are favored over the extremes (Wolf et al. 2007). Bird identity was modelled as a random effect on the intercept to account for pseudo-replication, and cohort to account for environmental stochasticity. We modelled fixed effects for sex (male, 1 or female, 0) and age (in calendar years) and age as a quadratic effect, to account for variation in fitness as explained by demography (Schroeder et al. 2012). We added sex as an interaction term with age to account for the extra-pair behaviour of older males (Girndt et al. 2018).

We modelled lifetime recruits and lifetime de-lifed fitness in the same way as the annual ones, but instead of age we used lifespan, or maximum age at year 2020. Because each bird was only represented once in this dataset, we only modelled cohort as a random effect.

We used Bayesian Markov Monte-Carlo simulations, using MCMCglmm (Hadfield 2010), to run all models. For all models we used inverse Wishart priors for random effects, and ran each over 343,000 iterations, with a burn-in of 3,000 and a thinning interval of 200. We visually checked the posterior trace plots for all model outputs and ensured that autocorrelation was below 0.1 and effective sample sizes between 1,000 and 2,000. The fixed effects were considered statistically significant when the 95% credible interval (CI) of its posterior distribution did not span zero.

### Null models and dominance interactions

We ran a node-permutation null-model by shuffling the identities of birds visiting the feeder between existing arrival times in our association matrices, thereby breaking any link between sociality and fitness, over 1000 randomized permutations (following Farine, 2017). We used these randomized association matrices to construct 1000 new networks and extracted the mode for our four centrality measures. We used these randomized centrality measures to re-run all fitness models. Finally, to exclude the possibility that dominance was interacting with our observed centrality measures, we tested for correlations between the centrality measures and dominance from videos recorded during the same period of our social network events. We represented individual dominance by calculating ELO ratings, based on antagonistic interactions at the bird feeder (for further details see Sánchez-Tójar 2018). We did not include the randomised centrality measures from our null models in these correlations.

## Results

The data consisted of 150 individual birds making 410,114 visits to the RFID feeder within our study period (mean = 2,734 visits per bird, SD = 8,116), across both events. Excluding birds that died prior to the start of our study or those that were ringed after, 160 tagged birds survived in our system in November 2015, an additional 90 birds were tagged prior to the 2017 breeding period, although not all survived to sampling. After constructing the arrival networks, we identified 3,783 associations between 118 PIT tagged individuals during 2015/2016, and 874 associations between 69 individuals in 2016/2017. These networks contained 66.3% of 122 and 26.3% of 205 breeding birds in 2016 and 2017 respectively. Combined, we had 130 records for annual and lifetime fitness from 102 individuals, 33 were recorded in both years (for summary statistics see Table 1). Degree and opposite-sex degree are closely correlated, implying that those with more opposite-sex associates also tend to have more associates of either sex (supplementary material table 1)

### Individual repeatability in sociality

We confirmed individual repeatability by week in all four centrality measures between 15 weeks in 2015/16: Degree, R=0.29 (0.15 – 0.39), Strength, R=0.22 (0.10 – 0.32), Centrality, 015 (0.03 – 0.27) and, Opposite sex degree, 0.27 (0.13 – 0.4); and, 13 weeks in 2016/17: Degree, R=0.29 (0.15 – 0.39), Strength, R=0.22 (0.10 – 0.32), Centrality, 015 (0.03 – 0.27) and, Opposite sex degree, 0.27 (0.13 – 0.4) (Table 1).

**Table 1.**
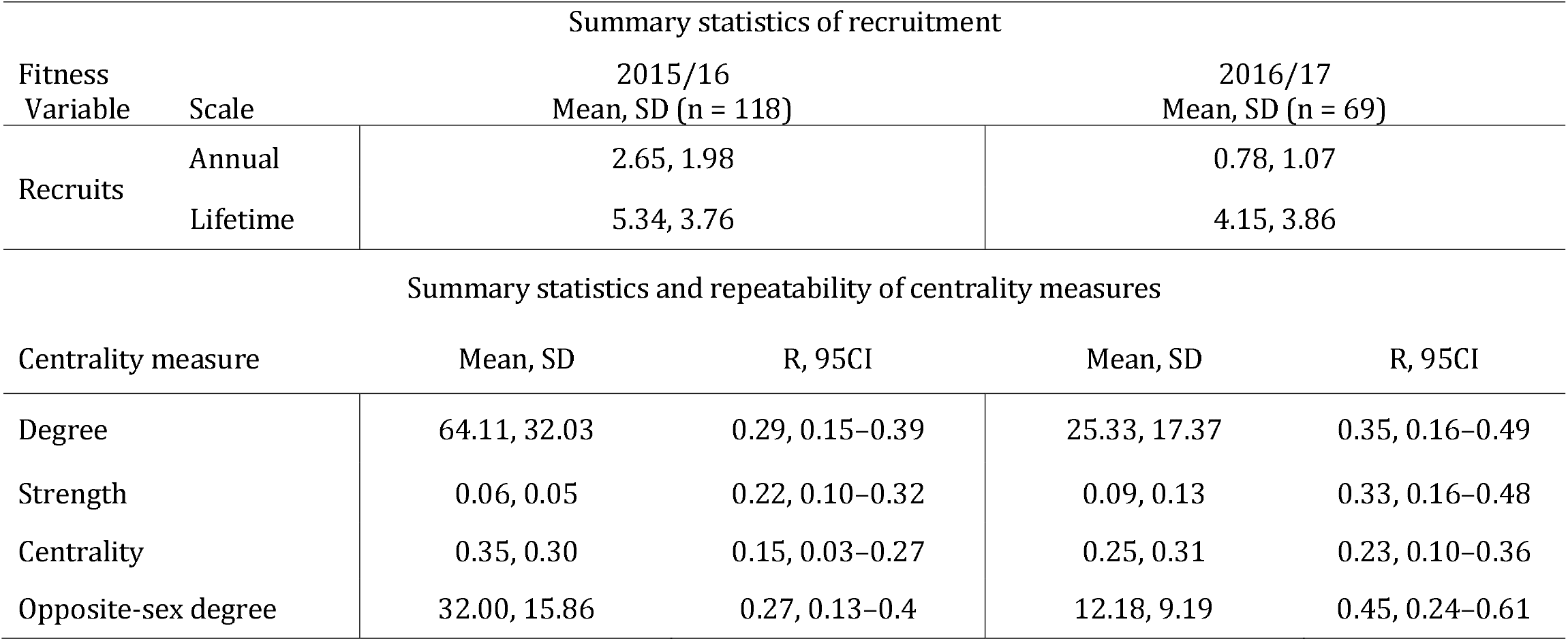
Summary statistics for recruitment and centrality measures for individual house sparrows on Lundy Island during two non-breeding events (November–January 2015/2016 and 2016/2017). Each measure is given as mean, standard deviation and sample size, and including repeatability (R) and 95% CI for centrality measures.

### Selection on sociality – annual and lifetime fitness

Opposite-sex degree had a statistically significant linear relationship with the number of annual recruits and annual de-lifed fitness. Strength and centrality had a negative quadratic association with annual recruitment (Figure 2; Table 2; Table 3). Age and sex both also predicted annual recruits, with younger individuals and females recruiting more offspring. Age also positively predicted annual de-lifed fitness (Table2; Table3).

**Table 2.** Annual recruitment model outputs from GLMMs for each of our four centrality measures (Degree, Strength, Centrality and Opposite-sex degree), derived of 410,114 visits to an RFID baited feeder by 150 individuals. Centrality measure of house sparrows on Lundy Island, modeled against annual recruits. We inferred significance where the 95% CI did not span zero, positive effects on the response variable are highlighted in red, and negative in blue.

| Annual recruits |  | Posterior mode<br>95% credible intervals |  |  |
| --- | --- | --- | --- | --- |
| Variable | Degree | Strength | Centrality | Opposite-sex degree |
| *Fixed terms* |  |  |  |  |
| (Intercept) | 3.49<br>1.46—6.11 | 3.69<br>1.58—6.46 | 4.19<br>1.60—6.31 | 3.78<br>1.58—6.50 |
| Centrality measure | 0.17<br>-0.01—0.34 | 0.22<br>-0.03—0.44 | 0.26<br>-0.01—0.45 | 0.25<br>0.05—0.41 |
| Centrality measure<sup>2</sup> | -0.1<br>-0.26—0.06 | -0.12<br>-0.32—0.07 | -0.16<br>-0.34—0.08 | -0.04<br>-0.16—0.10 |
| Sex (male) | -0.67<br>-1.41—0.05 | -0.7<br>-1.46—-0 | -0.74<br>-1.48—-0 | -0.57<br>-1.47—0 |
| Age | -1.14<br>-2.03—-0.30 | -1.15<br>-2.04—-0.28 | -1.27<br>-1.99—-0.26 | -1.07<br>-2.00—-0.26 |
| Age<sup>2</sup> | 0.05<br>-0.07—0.17 | 0.04<br>-0.08—0.17 | 0.06<br>-0.08—0.16 | 0.03<br>-0.07—0.17 |
| Age*Sex (male) | 0.21<br>-0.01—0.48 | 0.22<br>-0.02—0.46 | 0.28<br>-0.02—0.49 | 0.21<br>0—0.49 |
| *Random terms* |  |  |  |  |
| *Bird ID | 0<br>0—0.13 | 0<br>0—0.12 | 0<br>0—0.12 | 0<br>0—0.13 |
| *Cohort | 1.88<br>0.25—15.69 | 1.5<br>0.19—15.57 | 2.09<br>0—15.89 | 2.29<br>0.11—16.90 |
| Residuals | 0<br>0—0.27 | 0<br>0—0.28 | 0<br>0—0.29 | 0<br>0—0.24 |

**Table 3.** Lifetime recruitment model outputs from GLMMs for each of our four centrality measures (Degree, Strength, Centrality and Opposite-sex degree), derived of 410,114 visits to an RFID baited feeder by 150 individuals. Centrality measure of house sparrows on Lundy Island, modelled against lifetime recruits. We inferred significance where the 95% CI do not span zero, positive effects on the response variable are highlighted in red, and negative in blue. († Age in lifetime models denotes either lifespan, or age in 2020, whichever is greatest)

| Lifetime recruits |  | Posterior mode, 95% credible intervals (lower – upper) |  |  |
| --- | --- | --- | --- | --- |
| Variable | Degree | Strength | Centrality | Opposite-sex degree |
| *Fixed term* |  |  |  |  |
| (Intercept) | 0.85<br>0—1.93 | 0.85<br>0.02—1.95 | 0.99<br>-0.06—1.87 | 0.8<br>-0.03—1.83 |
| Centrality measure | 0.1<br>-0—0.25 | 0.1<br>-0.09—0.3 | 0.12<br>-0.07—0.31 | 0.14<br>-0.05—0.25 |
| Centrality measure<sup>2</sup> | -0.05<br>-0.19—0.11 | -0.01<br>-0.24—0.11 | -0.07<br>-0.24—0.11 | -0.03<br>-0.16—0.12 |
| Sex (male) | 0.33<br>-0.35—0.99 | 0.28<br>-0.36—0.91 | 0.28<br>-0.33—0.98 | 0.48<br>-0.24—1.03 |
| †Age | 0.1<br>0—0.23 | 0.11<br>0—0.24 | 0.09<br>0.01—0.25 | 0.12<br>0—0.24 |
| †Age*Sex (male) | -0.06<br>-0.20—0.07 | -0.07<br>-0.19—0.07 | -0.07<br>-0.2—0.06 | -0.06<br>-0.21—0.04 |
| *Random effects* |  |  |  |  |
| *Cohort | NA<br>0.07—2.78 | 0.47<br>0.07—2.76 | 0.45<br>0.06—3 | 0.39<br>0.10—2.77 |
| Residuals | NA<br>0—0.27 | 0.12<br>0.01—0.29 | 0.11<br>0.02—0.28 | 0.14<br>0—0.26 |

**Figure 2.** De-lifed fitness as response variables against Centrality measures from 8 linear mixed models, at two scales, from the Lundy Island house sparrows, derived of 410,114 visits to an RFID baited feeder by 150 individuals: Explanatory variables for Annual de-lifed fitness (A); and Lifetime de-lifed fitness (B), where ^2^ denotes a quadratic function, also shown in the four adjacent panels for A and B, and their 95% credible intervals. Credible intervals are given as solid bars for each explanatory variable, where a solid point denotes the posterior mode. Black bars denote no effect on the response variable; red denote a positive and blue, a negative, relationship with the response. In adjacent panels, quadratic functions of each response variable presented in A and B (on the Y axis: A Centrality, Degree, Opp. Degree, Strength, and B Centrality, Degree, Opp. Degree, Strength). Blue curves represent a negative interaction with fitness measures (on the X axis). Measures with no effect are not shown in figure. We found no link to sex, and age was also subject to stabilising selection, (given in supplementary material table 4).

None of our centrality measures statistically significantly predicted lifetime recruitment (Supplementary material table 4), however, all four had a statistically significant negative quadratic relationship with lifetime de-lifed fitness (Figure 2).

### Null models and dominance interactions

We found no link between fitness and sociality, nor any evidence of selection from our null models (supplementary material figure 3). Likewise, dominance was not strongly correlated with any centrality measure, implying that our method of assigning associations based on arrival time, rather than shared space at a bird feeder, is unlikely to be influenced by dominance (supplementary material table 1).

## Discussion

We found evidence for annual fitness benefits of sociality, where individuals with more opposite-sex associates had higher fitness in the breeding period, than those with fewer, but that this did not translate to lifetime fitness. For lifetime fitness, we found evidence for stabilising selection on sociality, including opposite-sex degree, suggesting that such benefits are only short-lived, or contextual, in a wild population.

We constructed our networks by linking dyads of birds that arrived together to a bird feeder, but ignored the time that they spent there, to eliminate most random associations. Other studies have adapted similar approaches in high-density and open feeder systems, or have considered the same implicit problems (Gomes, Boogert and Cardoso, 2021). Ferreira et al. (2020), for example, identified flocks arriving, but then defined associations by spatial proximity at a series of feeder boxes. Further research could optimize our approach for other systems, either by refining the time after which an individual is determined to have left the feeder (△_i_), or similarly, the time it takes for all members of a group to interact with the feeder upon arrival (△_t_). Further work may also consider defining associations only where a dyad visit together more often than would be expected by chance but doing so must also consider some method of retaining peripheral associations. Although RFID systems sample sociality well at a feeder, we cannot be sure that sociality traits are maintained in other contexts – future works might consider tracking social behaviour across time and space (For example see Ripperger and Carter 2021).

Where previous studies on wild birds have suggested links between aspects of sociality and annual reproductive success (for examples see Firth and Sheldon 2016; Kohn 2017; Beck, Farine, and Kempenaers 2020), we were also able to use lifetime measures, which better reflect the genetic contribution of the individual to population growth, and thus, fitness. This also allowed us to also describe how selection acts upon sociality across the population. We found that sociality had little influence on fitness at the annual scale, apart from for opposite-sex association, which was linked to increased recruitment and de-lifed fitness. Our study corroborates that annual fitness benefits described elsewhere, particularly regarding mate choice (Oh and Badyaev 2010; Beck, Farine, and Kempenaers 2021, Beck, Farine, and Kempenaers 2020) directly translate into increased annual fitness. At the lifetime scale, our study also provides some insight into the evolution of social behaviors, which we found to be maintained at the population average through stabilizing selection. We are therefore, to the best of our knowledge, the first study to link sociality with lifetime fitness benefits in a wild bird (but see Formica et al. 2021; Philson and Blumstein 2022). Our results may also suggest a mechanism for selection on sociality through enhanced mate choice, but the impact on survival was difficult to determine in this study. Sociality is predicted to increase survival through reduced predation risk or information transfer (Sorato et al. 2012; Hillemann et al 2020), but we found no evidence to suggest that either was selected for, through higher centrality, in our analyses.

Stabilizing selection in this case may be driven by factors such as high mate fidelity or changing sociality with age (Oh and Badyaev 2010; Albery et al. 2022), removing the need to constantly maintain opposite-sex associations over lifetime while maintaining individual fitness. However, those opposite-sex associations may also be beneficial in an extra-pair context from the male perspective (Beck, Farine, and Kempenaers 2020) and requires further research.

Our centrality measures were associated with lifetime, but not with annual de-lifed fitness, and only opposite-sex degree was associated with recruitment at the annual scale. We found no relationship between social centrality and dominance in our study using arrival time to define sociality, but aggressive interactions are probably also reduced over the non-breeding period (Summer-Smith 1963). None of our centrality metrics were linked with recruitment at the lifetime scales. Overall, de-lifed fitness better represents fitness as it is a relative measure of the contribution to population growth (Alif et al. 2022). The number of recruits, while intuitively appealing, is not relative, and in good years, more birds may have a higher number of recruits, while in poor years, having one recruit may be an achievement. As such, this measure is not always comparable between years and may explain our results. Further, recruitment is also dependent on parental effects and relationships within the breeding season, which were not quantified here, although they have been suggested elsewhere (Bebbington et al. 2017; Riehl and Strong 2018), whereas de-lifed fitness also captures long-term survival. We found that older males recruited more offspring, likely by virtue of older males siring more extra-pair offspring (Girndt et al. 2018). Likewise, younger birds had lower annual de-lifed fitness, because younger birds had not recruited any offspring in previous years that would contribute to their current de-lifed fitness.

In conclusion, we suggest a link between opposite-sex association and reproductive success at the annual scale, suggesting a mechanism for selection to shape social behaviour. At the lifetime scale we suggest that selection on sociality is stable, suggesting greater fitness for those at the population mean, in a wild population of passerine birds.

## Acknowledgments

We would like to thank the Lundy Landmark trust and the Lundy Field Society for their ongoing support, particularly Dean Jones, Rosie Ellis and Tom Carr. This research was supported by the QMEE CDT, funded by NERC grant number NE/P012345/1 (JD), a fellowship from the Volkswagen Foundation (JS), a grant from the German Research Foundation: Deutsche Forschungsgemeinschaft (JS), a European Research Council grant, CIG PCIG12-GA-2012-333096 (JS), and by NERC grant NE/J024597/1 (TB). We also thank two anonymous reviewers for their helpful feedback.

## Data availability

Analyses reported in this article can be reproduced using the data provided by Dunning, Jamie et al. (2022), Opposite-sex associations are linked with annual fitness, but sociality is stable over lifetime, Dryad, Dataset, https://doi.org/10.5061/dryad.z08kprrhd

