## Supplementary material for "Opposite-sex associations are linked with annual fitness, but sociality is stable over lifetime": Suppl files.

Figure 1. model outputs derived of a node-permutation null-model, randomizing bird identity against arrival time, without replacement, before constructing new networks and calculating mode network centrality measures from 1000 randomized permutations.

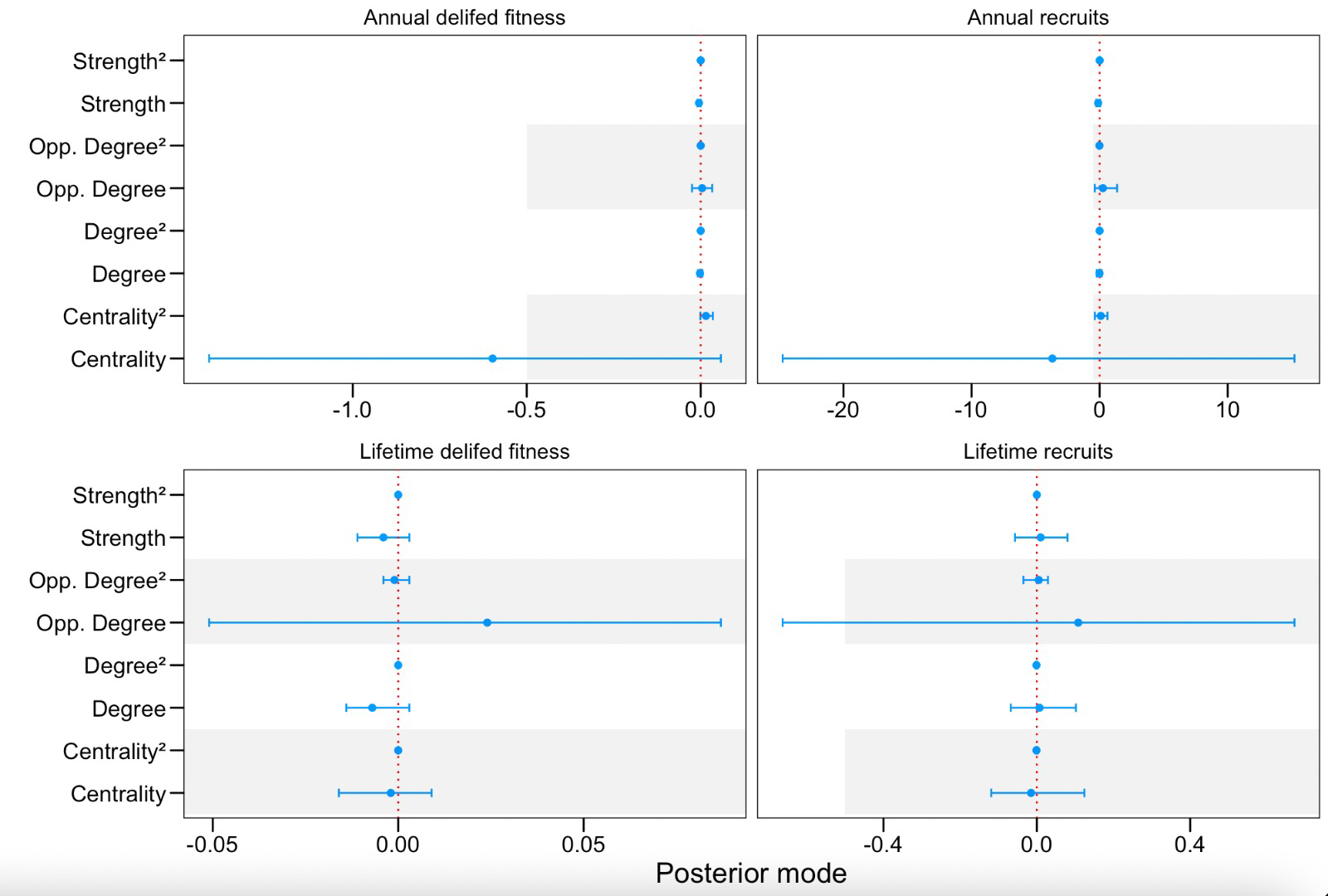

Figure 2. observed fitness as response variables against social measures from 8 linear mixed models, at two scales, from the Lundy Island house sparrows: Explanatory variables for Annual de-lifed fitness and Lifetime de-lifed fitness where ^2^ denotes a quadratic function and their 95% credible intervals. Credible intervals are given as solid bars for each explanatory variable, where a solid point denotes the posterior mode. Black bars denote no effect on the response variable; red denote a positive and blue, a negative, relationship with the response. In adjacent panels, quadratic functions of each response variable presented in A and B (on the Y axis: A Centrality, Degree, Opp. Degree, Strength, and B Centrality, Degree, Opp. Degree, Strength)

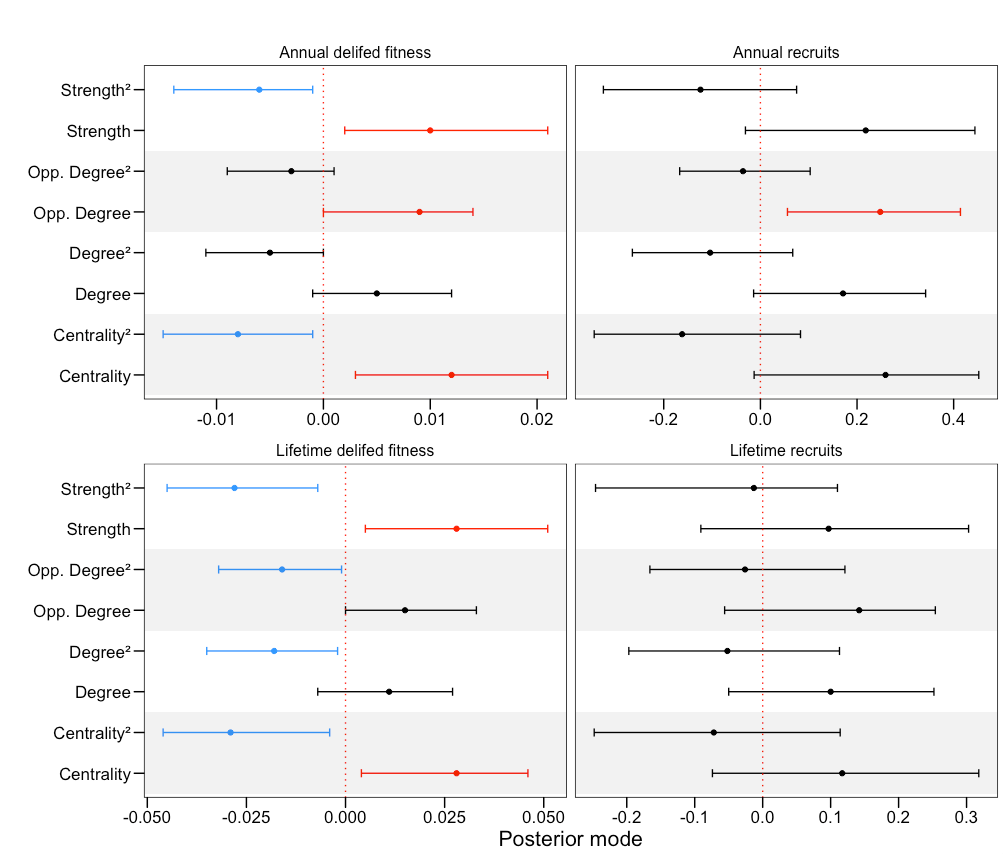

Table 1. showing the correlation between each social measure, but also same-sex degree and frequency of visit to the feeder.

Note that frequency is not strongly correlated with sociality measures. Also the three measures of degree are closely correlated, suggesting that sparrows with more associations of either sex, are also more likely to have more opposite sex friends, however, only opposite sex associations is linked with fitness – suggesting a role in mate choice.

|  | Degree | Centrality | Strength | Op_Degree | Same_Degree | Freq |
| --- | --- | --- | --- | --- | --- | --- |
| Degree | - | - | - | - | - | - |
| Centrality | 0.776 | - | - | - | - | - |
| Strength | 0.456 | 0.867 | - | - | - | - |
| Op_Degree | 0.962 | 0.753 | 0.434 | - | - | - |
| Same_Degree | 0.964 | 0.743 | 0.445 | 0.857 | - | - |
| Freq | 0.364 | 0.612 | 0.591 | 0.374 | 0.328 | - |
| ELO | 0.13 | 0.13 | 0.141 | 0.116 | 0134 | 0.131 |

Table 5. Model outputs from GLMMs for each of our four sociality measures (Degree, Strength, Centrality and Opposite-sex degree shown in Figure 2), against delifed fitness (Gaussian) at two scales in the Lundy house sparrow system. We inferred significance where the 95%CI do not span zero, positive effects on the response variable are highlighted in red, and negative in blue. Social variables are shown in Figure 2.

| Annual delifed fitness | | | Posterior mode, 95% credible intervals (lower – upper) | | | | |
| --- | --- | --- | --- | --- | --- | --- | --- |
| Variable | | Degree | Strength | | Centrality | Opposite sex degree | |
| *Fixed term* |  | |  | |  | |  |
| (Intercept) | -0.02 -0.05—0.03 | | -0.01 -0.04—0.03 | | -0.01 -0.04—0.03 | | -0.01 -0.05—0.03 |
| Sex (male) | -0.02 -0.04—0 | | -0.01 -0.04—0 | | -0.02 -0.04—0 | | -0.02 -0.04—0 |
| Age | 0.03 0—0.05 | | 0.03 0—0.04 | | 0.03 0—0.04 | | 0.03 0—0.05 |
| Age^2^ | 0 -0—-0 | | 0 -0—-0 | | 0 -0—-0 | | 0 -0—-0 |
| Age*Sex (male) | 0 -0—0.01 | | 0 -0—0.01 | | 0 -0—0.01 | | 0 -0—0.01 |
| *Random effects* |  | |  | |  | |  |
| *Bird ID | 0 0—0 | | 0 0—0 | | 0 0—0 | | 0 0—0.001 |
| *Cohort | 0 0—0 | | 0 0—0 | | 0 0—0 | | 0 0—0.002 |
| Residuals | 0 0—0 | | 0 0—0 | | 0 0—0 | | 0 0—0.001 |
| Lifetime delifed fitness | | | | Posterior mode, 95% credible intervals (lower – upper) | | | |
| Variable | Degree | | Strength | | Centrality | Opposite sex degree | |
| *Fixed term* |  | |  | |  |  | |
| (Intercept) | 0.02 -0.05—0.07 | | 0.01 -0.05—0.07 | | 0.01 -0.05—0.08 | -0.02 -0.06—0.05 | |
| Sex (male) | 0.03 -0.04—0.09 | | 0.03 -0.05—0.08 | | 0.03 -0.03—0.08 | 0.02 -0.03—0.09 | |
| †Age | 0.01 -0—0.02 | | 0.01 -0—0.01 | | 0.01 -0—0.01 | 0.01 -0—0.02 | |
| †Age*Sex (male) | -0.01 -0.02—0 | | 0 -0.01—0.01 | | 0 -0.01—0 | -0.01 -0.02—0 | |
| *Random effects* |  | |  | |  |  | |
| *Cohort | 0 0—0 | | NA 0—0 | | 0 0—0 | 0 0—0 | |
| Residuals | 0 0—0 | | NA 0—0 | | 0 0—0 | 0 0—0 | |
